# Aperiodic EEG Activity Provides a Linear, Bidirectional, and Spatially Uniform Marker of Subjective and Objective Vigilance in Humans, Both Within and Across States

**DOI:** 10.1101/2025.08.30.673229

**Authors:** Henry Hebron, Aurelie M. Stephan, Jacinthe Cataldi, Francesca Siclari

## Abstract

Vigilance is increasingly conceived as a continuum, ranging from full alertness to deep sleep. Despite its fundamental role in cognition, behaviour, and health, reliable physiological markers of vigilance remain limited, and clinical assessments often rely on subjective or time-consuming evaluations. Traditionally, vigilance has been estimated through visual inspection of the electroencephalogram (EEG), identifying recognizable oscillatory patterns like rapid, wakefulness-defining alpha waves (∼10 Hz) and large slow waves (∼1 Hz) which typify sleep. However, these oscillatory features often appear only intermittently and follow complex, non-linear trajectories across time, space, and frequency, limiting their utility for automated, continuous tracking of vigilance.

Recent research has shifted attention to the non-oscillatory, or *aperiodic*, component of the EEG, which may follow simpler dynamics and offer a more robust index of brain state. Yet most studies often continue to use narrowly defined, discrete vigilance states and transitions in only one direction (e.g., from wakefulness to sleep), without jointly examining oscillatory and aperiodic activity.

Here, we address these key gaps by evaluating the capacity of both oscillatory and aperiodic features of EEG power spectra, derived from high-density recordings, to predict vigilance as a continuous variable. Across three independent datasets, we consistently show that although oscillatory features reliably track changes in vigilance, they are unequivocally outperformed by aperiodic activity. Aperiodic features demonstrate a stronger, more linear, and spatially consistent relationship with both objective and subjective indices of vigilance, offering a more robust and scalable physiological marker of this fundamental feature of the brain.

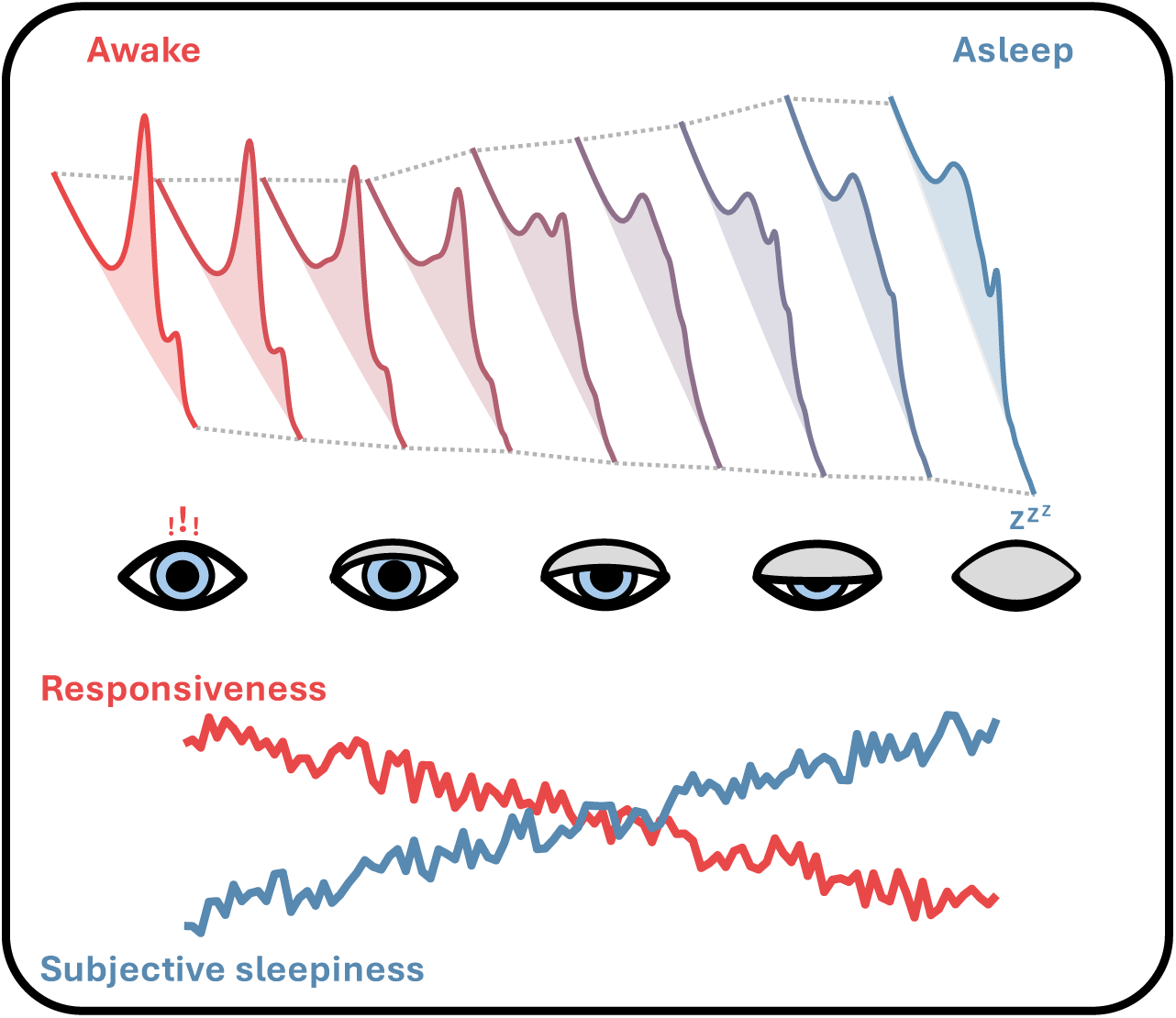

## 2 INTRODUCTION

Life depends on repeated, reversible, and reliable transitions between wakefulness and sleep [1,2]. The space between these two fundamental states is increasingly conceived as a *continuum* of vigilance [1–5], beginning with extreme alertness, and progressing through drowsiness, shallow sleep, deep sleep, and beyond.

The brain’s precise positioning on this continuum has important clinical implications, since imbalances in vigilance can severely disrupt everyday functioning. Individuals with impaired vigilance, such as those with hypersomnia or related disorders, may struggle to remain alert (for review, see [6]), while others with excessive vigilance – like hyperarousal in insomnia [7] – find it difficult to initiate or maintain sleep. Both conditions are associated with poorer health outcomes and increased disease risk [8–11]. Altered vigilance is also a feature of many neurodegenerative disorders and may even contribute to their progression [12,13]. More broadly, fluctuations in vigilance, whether due to a sleep disorder or as a result of pathology, represent a widespread public health concern with significant societal costs [14,15], particularly due to reduced workplace productivity [11,14,15] and increased risk of traffic and other accidents [15,16].

Beyond its clinical and societal relevance, vigilance is a critical factor in virtually all investigations of the brain, modulating perception, cognition, attention, and physiology [1,2,4,17–20]. Tracking vigilance, therefore, is imperative for interpreting neural and behavioural data [20]. However, robust and easily interpretable physiological measures of vigilance remain elusive.

Traditional approaches rely on expert visual evaluation of the electroencephalogram (EEG) to classify vigilance into discrete stages, typically following systems like the AASM [21] or Hori [22,23] scoring criteria, based on the presence of characteristic neural oscillations. Alpha rhythms (8–12 Hz) are used to mark wakefulness, while slow oscillations and spindles typify non-REM sleep. Clinical tools such as the Maintenance of Wakefulness Test (MWT [24,25]) and the Multiple Sleep Latency Test (MSLT [25,26]) rely on such EEG-based markers of sleep onset. However, these clinical procedures are labour-intensive, time-consuming for both the patient and the clinician, and subject to interpretative variability, limiting their scalability and objectivity [24,27,28] . These measures are also dependent on motivation [24], and do not always correlate well with subjective experience of vigilance [6,29–31]. In addition, these conventional approaches capture only coarse, 30 second epoch-based changes [5,27] and primarily index sleep propensity rather than drowsiness [29], limiting their ability to describe the vigilance continuum[5] and raising concerns about their use in safety-critical contexts such as driving or workplace assessments [6].

Although this traditional oscillation-centric perspective on vigilance could, in principle, be refined by improving the methods by which the brain’s rhythms are quantified, a greater challenge, still, lies in the inherent complexity of the oscillatory dynamics themselves. Oscillatory features often assume non-linear trajectories across time, frequency, and brain regions with changing vigilance [1–4,32,33]. For instance, oscillations of between 8 and 12 Hz become rapidly diminished with the transition to light sleep in posterior regions of the cortex (alpha oscillations) before slowly emerging in more anterior areas (slow sleep spindles) as sleep deepens [4,33].

These issues are compounded further by the techniques most commonly used to *quantify* the rhythms of the brain – frequency-limited band power (i.e., the summation of energy derived from Fourier decomposition, within predefined frequency bands, such as delta, theta, alpha, beta, or gamma). Band power, whilst often treated as equivocal with oscillatory activity, actually represents a mixture of rhythmic (periodic) and arrhythmic (aperiodic) activity [34–39]. Moreover, since brain rhythms are often non-sinusoidal, non-stationary, and intermittently absent, they frequently violate the assumptions underlying conventional spectral methods such as Fourier analysis.

To address these limitations, recent studies have adopted approaches that explicitly separate periodic and aperiodic components of the EEG power spectrum [34–50]. Oscillatory activity is modelled as (gaussian) peaks superimposed on a 1/f exponential decay-like background, representing aperiodic neural activity. Some characteristic oscillations which are captured by the periodic component include, but are not limited to, alpha and theta rhythms in wakefulness, and sleep spindles in non-REM sleep. The aperiodic component, meanwhile, captures a more general measure of the relative power of high and lower frequencies across the full spectrum.

This decomposition has not only improved the precision of oscillatory analyses [34–39] but has also revealed the aperiodic signal itself to be a powerful marker of brain state, capable of distinguishing sleep stages [45–50], tracking depth of anaesthesia [41,45], and reflecting broader neurophysiological processes [39,40,44,48,49,51–55].

Despite this progress, key limitations remain. Many studies to date have continued to focus on classifying discrete sleep stages, rather than modelling vigilance as a continuous variable. Few have incorporated both subjective (e.g., self-report) and objective (e.g., reaction time, responsiveness) measures of vigilance, and many rely on either periodic or aperiodic components in isolation, neglecting the possibility that each may offer unique and complementary insights. Finally, much of the existing work has been conducted using low-density EEG, limiting the ability to capture the full spatial structure of vigilance-related neural activity.

In the present work, we aim to resolve these limitations by systematically comparing the contributions of periodic and aperiodic EEG features to continuous measures of vigilance, both subjective and objective, using high-density EEG recordings. Across three independent studies, we track neural and behavioural fluctuations within and across states, including transitions between wakefulness and non-REM sleep in both directions. Our goal is to identify a physiologically grounded, spatially consistent, and temporally sensitive marker of vigilance that generalises across brain states.

With a striking degree of consistency across the datasets, we find that while both periodic and aperiodic EEG features capture changes in vigilance, they exhibit distinct spatial and temporal profiles, suggesting they are representing different elements of brain state. Critically, aperiodic activity consistently outperforms oscillatory measures, varying in a simple, linear, and spatially global manner with changes in vigilance. These findings support the use of aperiodic EEG features as a robust physiological analogue of vigilance and offer a path toward more objective and scalable tools for both clinical assessment and neuroscience research.

## 3 RESULTS

Vigilance is an intuitive concept – we all know how it *feels* to be alert or drowsy. But it may be difficult to explicitly and quantitatively define. Here, to cover all bases, we took a multi-faceted approach to vigilance, exploring several complementary aspects (**Figure 1**). Firstly, we assumed an *<u>agnostic</u>* perspective, simply considering that vigilance decreases with time during the first portions of nocturnal sleep (**3.1**). Secondly, we explored a *<u>subjective</u>* metric of vigilance, as quantified by questions about feelings of sleepiness, immediately following awakening in a serial awakening paradigm [31,56,57] (**3.2**). Finally, we implemented an *<u>objective</u>* measure of vigilance, analysing reaction times and responsiveness during a lengthy button-press task, in which participants drifted between wake and sleep [20] (**3.3**). Across each of these approaches, we related the degree of vigilance to both aperiodic and oscillatory components of the neural power spectrum, as derived from high-density EEG recordings, using the SPRiNT toolbox [36]. In particular, the exponent of the aperiodic component and the Signal:Noise ratio (SNR – i.e., height of fitted gaussian, indicative of magnitude of oscillation) and centre-frequency of the periodic component were explored.

**Figure 1.**
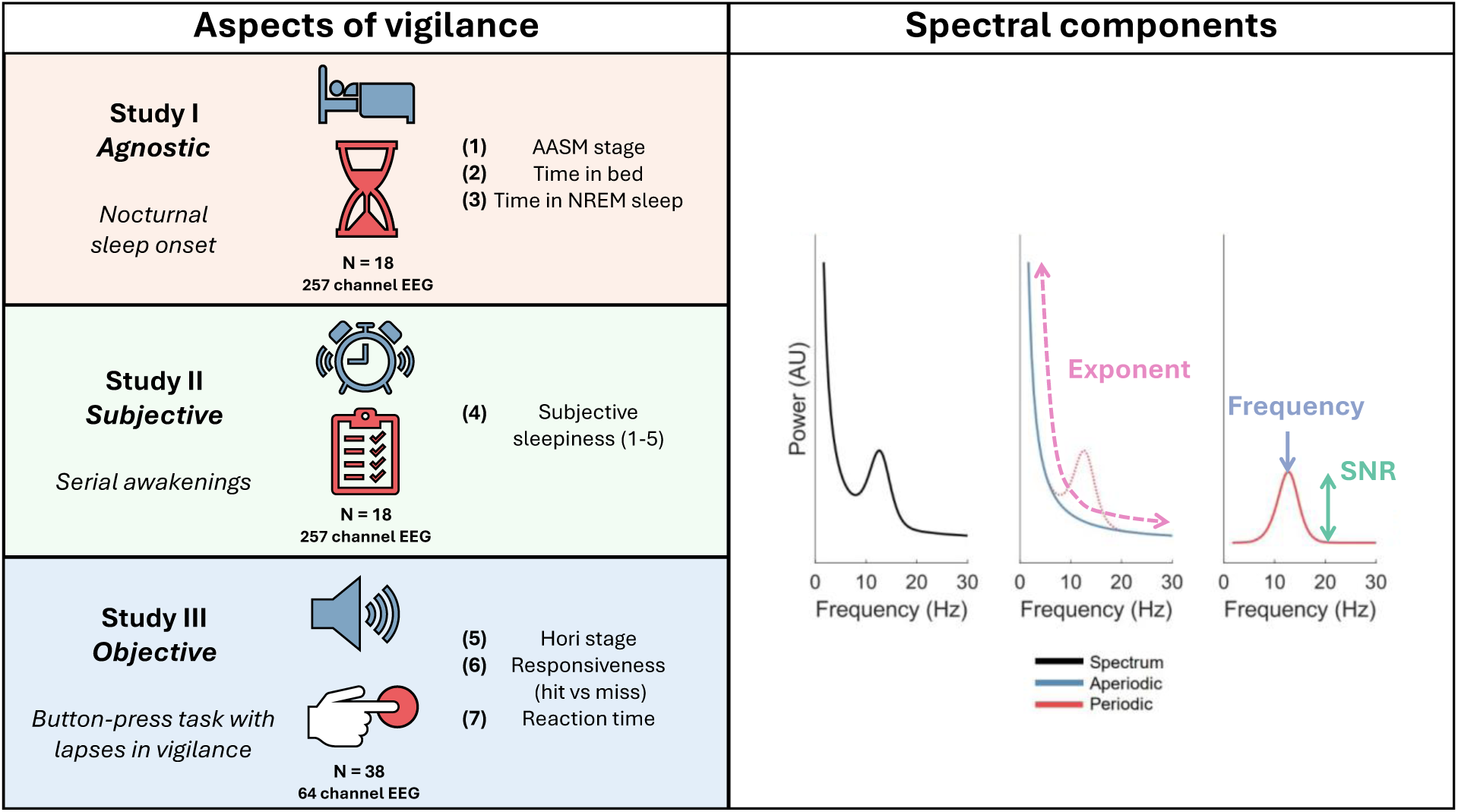
Overview of experimental approach to defining vigilance, and extracting physiological measures. Shown on the left are the three datasets utilised, complete with their accompanying aspects of vigilance (1)-(7), sample sizes, and EEG channel number. Shown on the right are the periodic and aperiodic parameters derived from neural power spectra – exponent, centre frequency and signal:noise ratio (SNR) - demonstrated using a synthetic, idealised, power spectrum.

### 3.1 Study 1 – Agnostic

#### Aperiodic and Periodic Components Both Distinguish Sleep Stage, but with Different Topographical Profiles

Although our intention was to arrive at a *continuous* physiological analogue of vigilance, it was important to first replicate previous investigations of traditional *categorical* definitions of vigilance. Hence, we first sought to derive the relationship between AASM-defined stages [21] (W, N1, N2, N3) and aperiodic and periodic spectral components, initially focusing on those oscillations which characterise wakefulness and sleep onset (theta and alpha) (**Figure 2**). Whilst spectral exponent was steepest in frontal-medial channels across sleep stages (**Figure 2B**), a strong positive effect of stage was observed across *all* channels (**Figure 2C**) – i.e., a steeping slope with the progression from wake to N3. The relationship between oscillatory components and stage, meanwhile, was more complex, showing divergent dynamics across the scalp, with anterior theta SNR decreasing and posterior theta SNR increasing (**Figure 2E**), and alpha SNR following the precise opposite trajectory (**Figure 2F**). It may be the case that this increase in anterior alpha SNR is capturing the emergence slow sleep spindles (most prominent in frontal regions [58–61] and increasingly prominent with the progression of NREM sleep [58,61]) – and overlapping in frequency with alpha oscillations) already demonstrating the complications of capturing vigilance on the basis of specific frequency bands.

**Figure 2.**
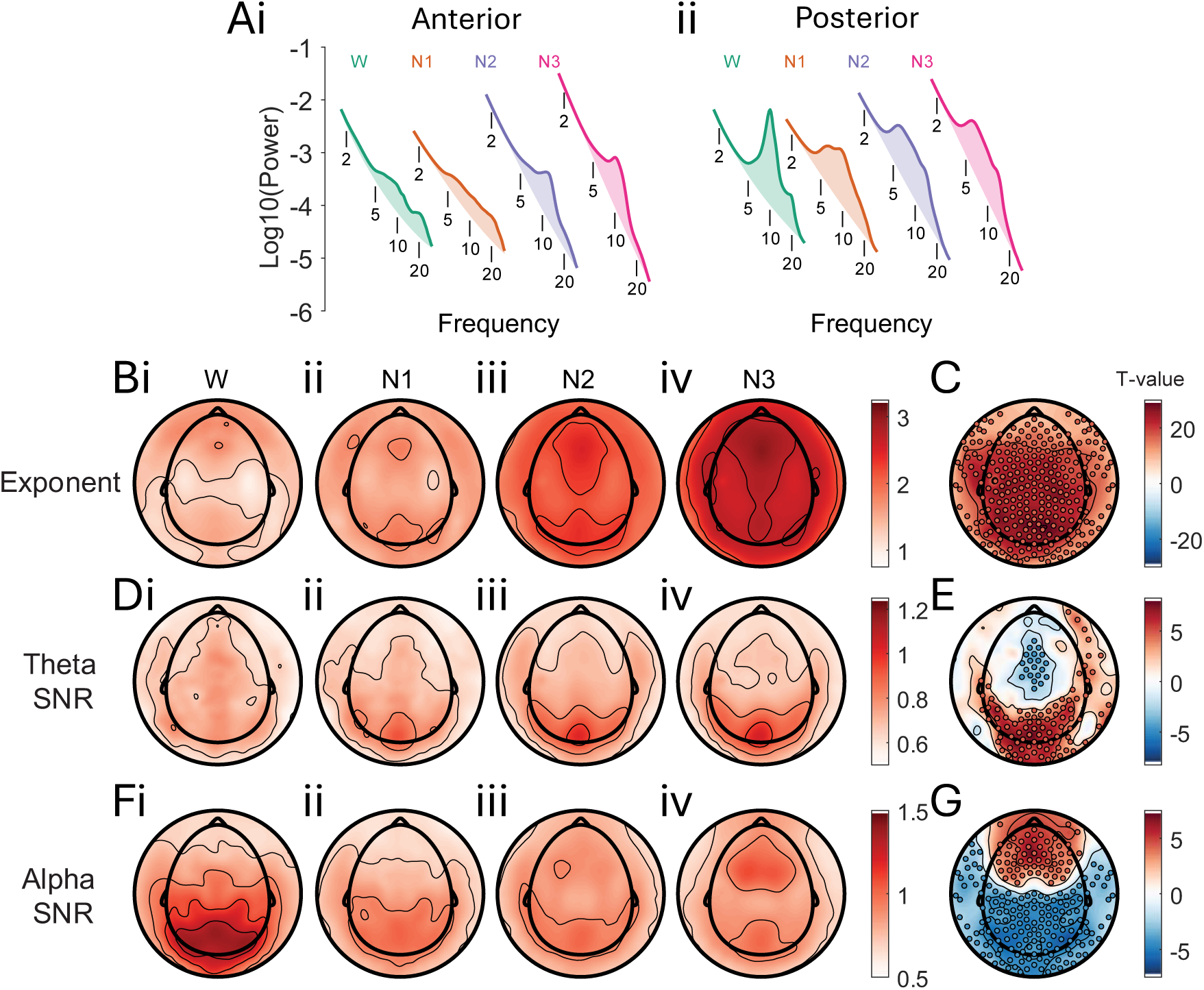
Aperiodic and periodic spectral features over discrete vigilance states. **(A)** Power spectra across wakefulness (W), and the three stages of NREM sleep (N1, N2, N3) at **(i)** anterior and **(ii)** posterior electrode clusters (see methods); **(B)** Topographic representations of spectral exponent in **(i)** wakefulness (W) , **(ii)** NREM1 (N1) , **(iii)** NREM2 (N2), and **(iv)** NREM3; **(C)** Topographic representation of effect of AASM-defined stage (W, N1, N2, N3) on exponent as per linear mixed effects model [exponent ∼ vigilance state + (1|participant)]; **(D)(E)** same as **(B)** and **(C)** but for theta SNR; )]; **(F)(G)** same as **(B)** and **(C)** but for alpha SNR. Across all topographies, red scatter points indicate a (FDR-corrected statistically significant, p<.05) positive relationship between vigilance state and the EEG feature and blue scatter points indicate a (FDR-corrected statistically significant, p<.05) negative relationship. Exponent units are a.u. Hz^-1^, and SNR units are a.u.

#### Aperiodic and Periodic Components Show Distinct Spatio-Temporal Trajectories During the Transition to Sleep

In order to take a step further than the traditional discretisation of brain state, we plotted our metrics of interest across the first hour of the night, following lights off, uncovering their dynamics with decreasing vigilance (**Figure 3**). A fairly linear dynamic was observed in the spectral exponent across time, at both anterior and posterior regions, an effect which, once again, could be observed with a similarly strong intensity at all channels (**Figure 3A**). Theta and Alpha SNR, however, exhibited more complex dynamics in space and time, with posterior SNR theta and alpha changing in opposite directions in the first 10 minutes before levelling out and anterior SNR changing more gradually across the hour (**Figure 3 Bi**, **3Ci**). This effect of time on oscillations, once more, had a similarly complex topographical identity which mirrored the one previously shown for of AASM-defined stage (**Figure 3Bii, 3Cii**).

**Figure 3.**
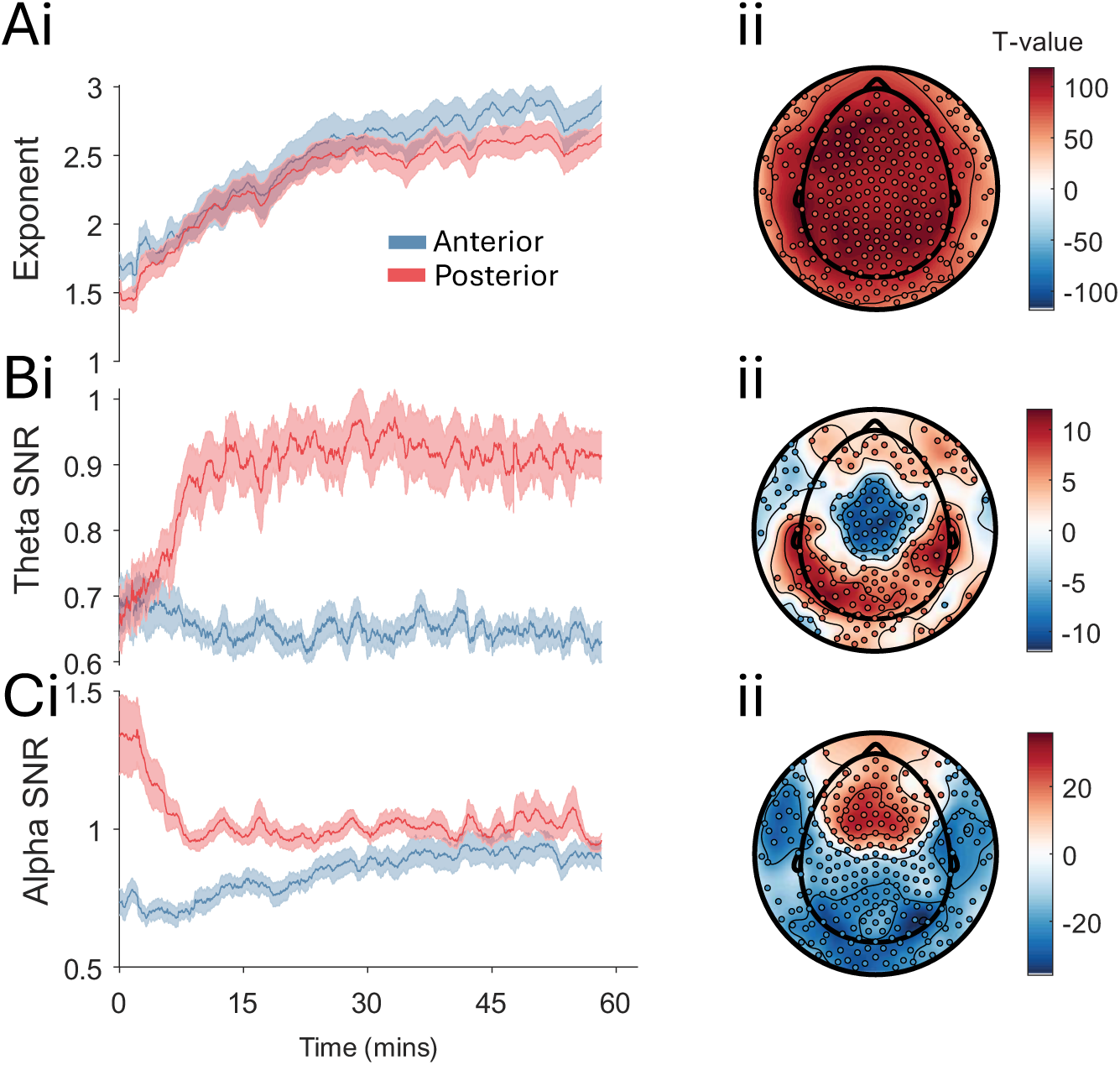
Aperiodic and periodic spectral features over time. **(A)(i)** Time-series of exponent across the first hour of the night recording (with 0 corresponding to lights-off, the beginning of the sleep opportunity) at anterior and posterior electrode clusters (see methods), **(ii)** Topographic representation of effect of time on exponent as per linear mixed effects model [exponent ∼ time + (1|participant)]; **(B)** Same as **(A)**, but for theta SNR; **(C)** Same as **(A)**, but for alpha SNR. Across all topographies, red scatter points indicate a (FDR-corrected statistically significant, p<.05) positive relationship between time and the EEG feature and blue scatter points indicate a (FDR-corrected statistically significant, p<.05) negative relationship. Data were smoothed using a 50-window (76.5 seconds) moving mean Exponent units are a.u. Hz^-1^, and SNR units are a.u.

#### Aperiodic and Periodic Components Track Vigilance *Within* Sleep

Since we are interested in vigilance as a continuum, in addition to contrasting *between* or *across* discrete states, it must be mappable in a continuous manner *within* a behavioural state. This is vital for a measure to be more useful than traditional sleep-scoring. To investigate this, we analysed the effect of time *within* NREM sleep on aperiodic and periodic spectral components in the first 10 minutes of N2 and N3 sleep separately, once more operating under the assumption that vigilance decreases with time (**Figure 4**). Since the focus was exclusively on NREM sleep, we explored oscillations in the spindle frequency range, whilst the aperiodic component continued to be analysed as before – i.e., the spectral slope.

**Figure 4.**
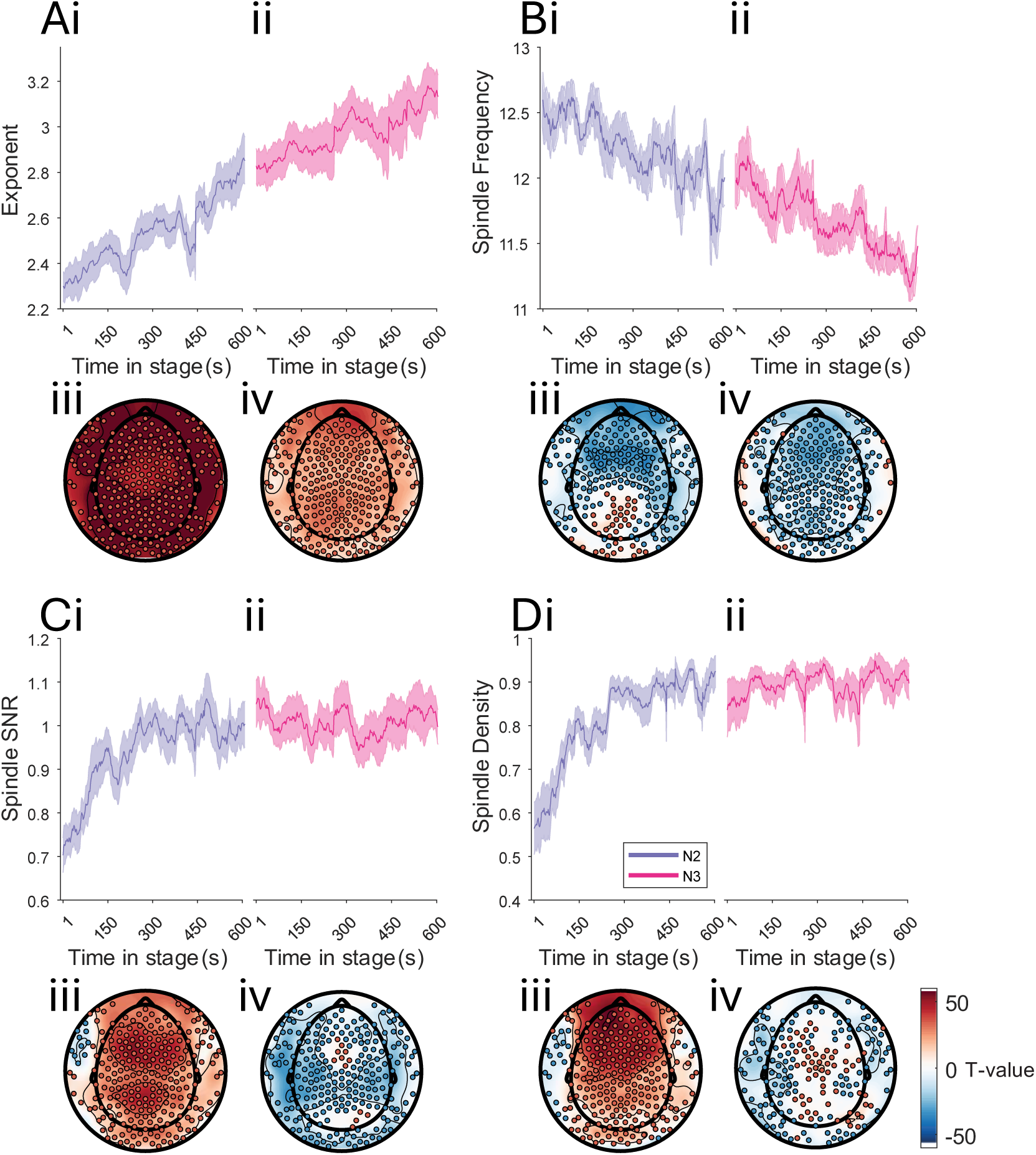
Periodic and aperiodic spectral features within NREM sleep. **(A)** Time-series of exponent across the first contiguous ten minutes of **(i)** NREM2 and **(ii)** NREM3 at Fz, **(ii)** Topographic representation of effect of time in **(iii)** NREM2 and **(iv)** NREM3 on exponent as per linear mixed effects model [exponent ∼ time_in_NREM + (1|participant)]; **(B)** Same as **(A)**, but for spindle frequency; **(C)** Same as **(A)**, but for spindle SNR. **(D)** Same as **(A)**, but for spindle density. Across all topographies, red scatter points indicate a (FDR-corrected statistically significant, p<.05) positive relationship between time and the EEG feature and blue scatter points indicate a (FDR-corrected statistically significant, p<.05) negative relationship – FDR-corrected *p*<.05. Data were smoothed using a 25-window (39 seconds) moving mean. Exponent units are a.u. Hz^-1^, and SNR units are a.u.

Once more, the spectral slope was observed steepening with time, within both N2 (**Figure Ai**) and N3 (**Figure 3Aii**) sleep – albeit, starting at an expectedly steeper point with the beginnings of N3 sleep. This effect was spatially global, as before, but with a stronger linear relationship between time and spectral slope in N2 (**Figure 3Aiii**) than N3 sleep (**Figure 3Aiv**). The features of oscillations in the sleep spindle range, in contrast, displayed a more complex dynamic across their different features. Whilst their frequency decreased in a fairly coherent and spatially global manner across the 10 minute episode (**Figure 3B**), their SNR and density showed a more complex dynamic, strongly increasing in the first 5 minutes of N2 sleep (**Figure 3Ci**, **Di**), before plateauing before the onset of N3 sleep (**Figure 3Cii**, **3Dii**). The topography of these effects showed a spatially-broad increase with time in N2 sleep (**Figure 3Ciii**, **3Diii**), which, unlike the exponent and the spindle frequency, was not maintained into N3 sleep (**Figure 3Civ**, **3Div**). These within-oscillation nuances suggested that spindles may serve as a less useful or ubiquitous analogue of vigilance, and may follow more complex dynamics.

In summary, the aperiodic spectral component appears to provide a more encompassing measure of vigilance than its oscillatory counterparts – more straightforward in both time and space and across states.

### 3.2 Study 2 - Subjective

Whilst our first analyses provided valuable insights into which measures might best capture vigilance, this assumption of a decrease in vigilance with time is simplistic, and may be limited in its utility. Hence, we sought to relate our established metrics of interest with more meaningful readouts of vigilance, which brought us to the subjective experience of sleepiness, measured by questionnaire following serial-awakenings from NREM sleep[31,57].

#### The Trajectory of Spectral Components During Awakenings Mirrors that of Sleep Onset, but Only the Aperiodic Component Robustly Predicts Subjective Sleepiness

First, we again explored the dynamics of spectral exponent, theta SNR, and alpha SNR across time, but this time, during a very sharp *increase* in vigilance – upon awakening from NREM sleep following an alarm. We then related the EEG measures, as averaged over the last 2 minutes of sleep and the 10 seconds following awakening separately, to post-awakening subjective reports of sleepiness.

The spectral slope showed a pronounced drop in its steepness following the awakening, arriving at the same point in both anterior and posterior regions, despite beginning higher in the anterior region (**Figure 4Ai**). Spectral exponent showed a largely global relationship between subjective sleepiness when calculated prior to the awakening (i.e., a steeper spectral slope was related to feelings of greater sleepiness, **Figure 4Aii**). This relationship was captured by *every* channel when calculated following the awakening (**Figure 4Aiii**), consistent with the expectations established in **Study 1**.

Although theta and alpha SNR also met these expectations in their dynamics across the awakening in the posterior region (**Figure 4Bi**, decrease in theta, **4Ci** increase in alpha), they captured subjective vigilance poorly, both before and after awakening (**Figure 4B**, **4C**), with only a few channels showing a statistically significant relationship with subsequent sleepiness, when measured before the awakening.

Once more, the aperiodic spectral component provided a stronger and more straightforward readout of vigilance than the oscillatory component across states.

#### Objective

Having related spectral features to our agnostic assumptions and subjective aspects of vigilance, we sought to derive a more objective measure – a behavioural readout that could both describe vigilance in a binary between-states manner (responsive vs unresponsive) and in a continuous within-states manner (speed of responsiveness). To this end, we deployed our analyses on a previously collected auditory-masking task dataset, a simple task in which participants spent 2 hours responding to sound stimuli by pressing a button. Owing to the length of the task, and the conditions under which it was performed (dark room, eyes-closed), participants exhibited numerous drifts and lapses in vigilance, providing the perfect substrate for an objective definition.

#### Hori Stages Align well with Previously Established Physiological Readouts of Vigilance

In this dataset, EEG traces were once again categorised into pre-defined discrete states of vigilance. However, in this case the Hori system was used, a set of rules derived to better describe the continuum from wakefulness into shallow sleep, in which vigilance is divided into 9 different states. When comparing Hori stages (as with the AASM sleep stage classifications, time in bed, and subjective sleepiness) the same dynamics were once more observed – a global steepening of the spectral slope, accompanied by a decrease in posterior alpha, and an increase in posterior theta (**Figure** ).

#### Spectral Components Predict Responsiveness and its Speed, Consistent with Previous Analyses

We then compared the EEG in the moments before hits and misses (i.e., responsive vs unresponsive) – using the 4 seconds of EEG before the stimulus was presented. The same physiological features which separates vigilance state once again also distinguished these behavioural states (**Figure 7A, 7B, 7C**), i.e., global spectral exponent, posterior theta SNR, and posterior alpha SNR. The relationship was strongest for the exponent, as before.

**Figure 5.**
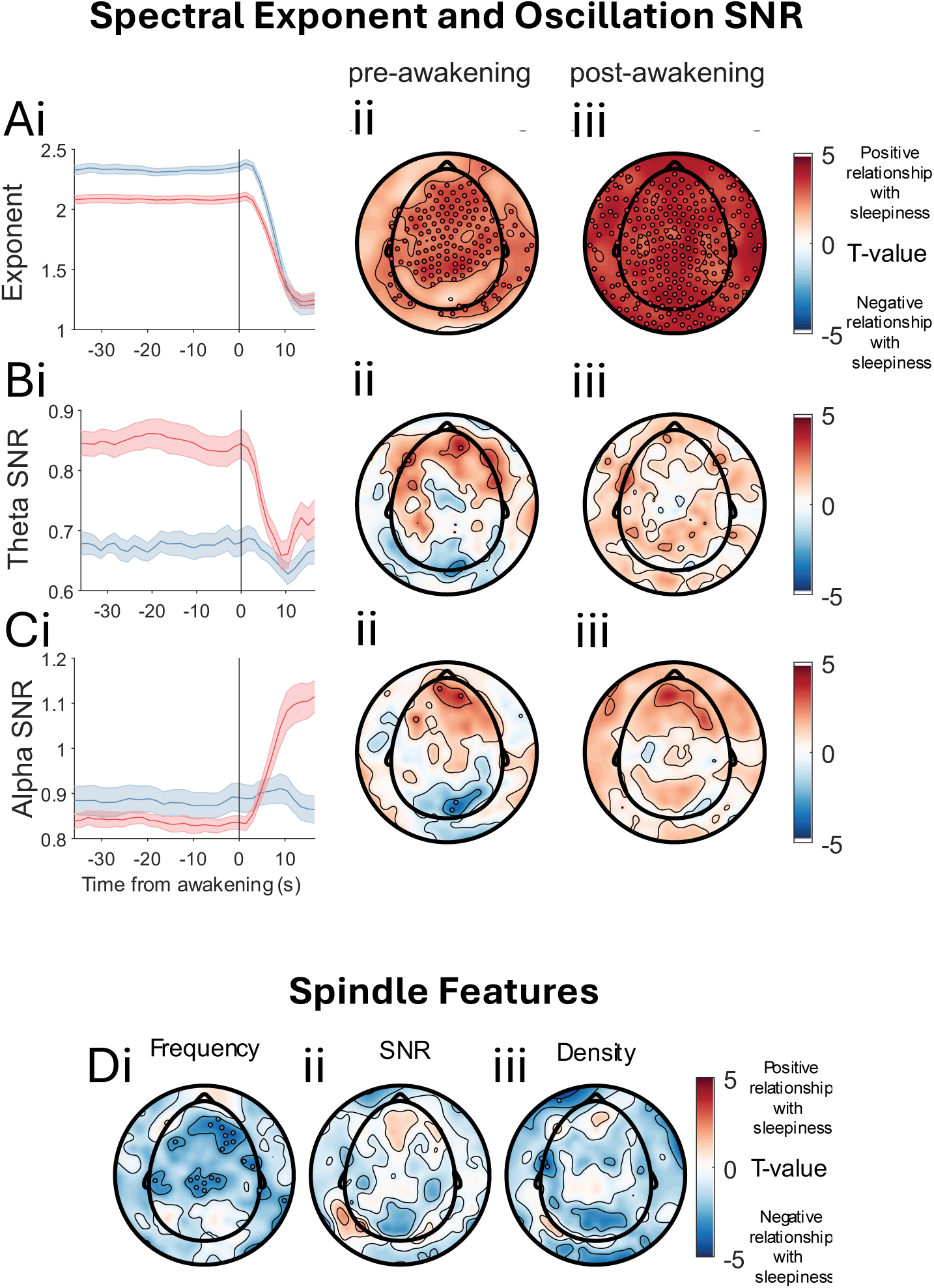
The Trajectory of Aperiodic and Oscillatory Features During Awakenings, and Their Ability to Predict Subsequent Subjective Sleepiness. **(A) (i)** time series of exponent at an anterior (blue) and posterior (red) channel in the moments preceding (30 seconds) and proceeding (15 seconds) awakenings from NREM sleep. Shaded error bars indicate standard error of the mean; **(ii)** topographical representation of relationship between exponent averaged across the two minutes preceding awakenings and subjective sleepiness following awakening; **(iii)** same as **(Aii)**, but for 10 seconds proceeding awakening. **(B)** same as **(A)** but for theta SNR. **(C)** same as **(A)**, but for alpha SNR. **(D)** relationship between subjective sleepiness and pre-awakening spindle **(i)** frequency, **(ii)** SNR, and **(iii)** density. Red scatter points on topographies indicate a (FDR-corrected statistically significant, p<.05) positive relationship between subjective sleepiness and the EEG feature in question, whilst blue scatter points indicate a (FDR-corrected statistically significant, p<.05) negative relationship. Data were smoothed using a 4-window (7.5 seconds) moving mean. Exponent units are a.u. Hz^-1^, and SNR units are a.u.

**Figure 6.**
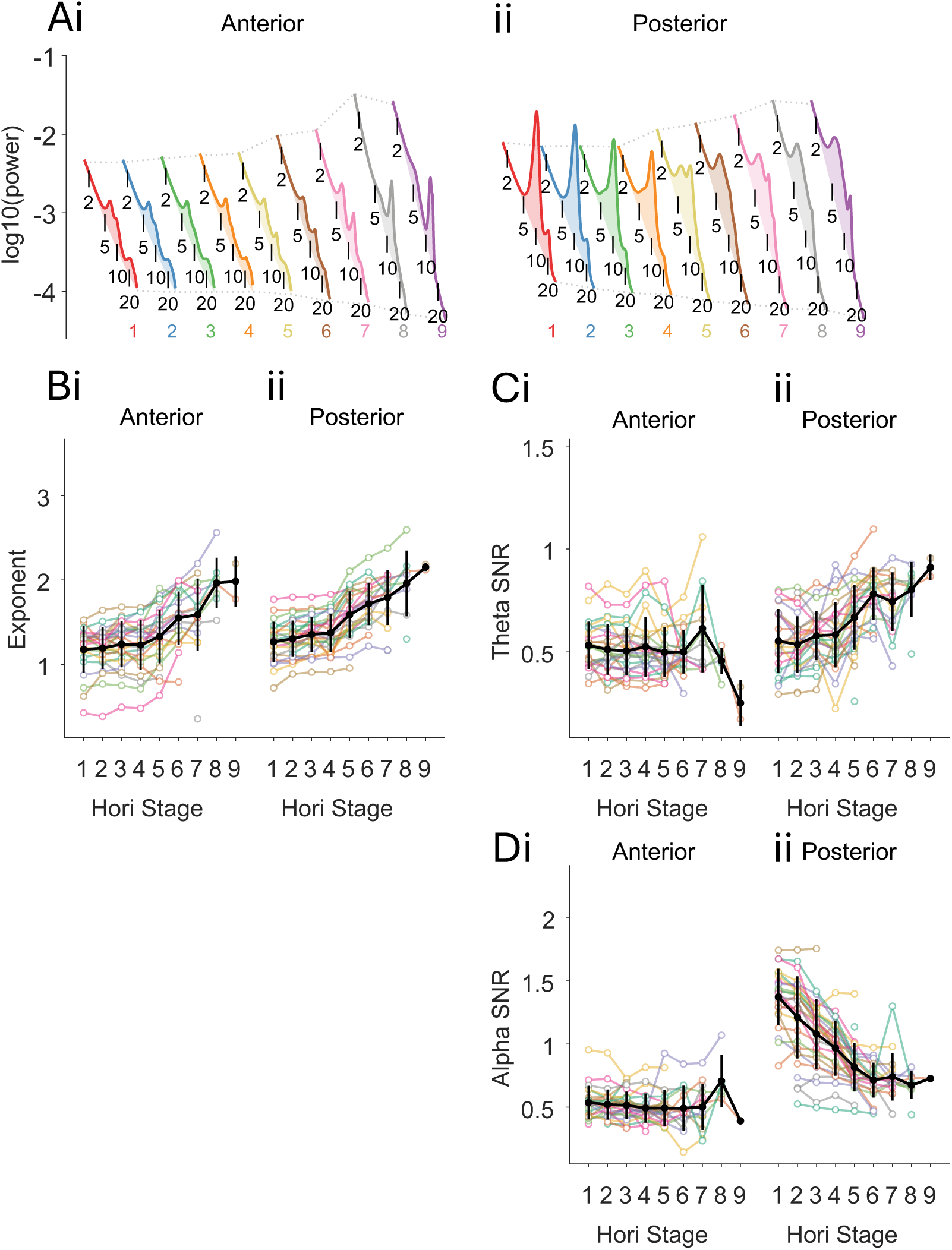
Aperiodic and Oscillatory Features Across Hori Stages of Vigilance. **(A)** Power spectra across Hori stages at **(i)** an anterior and **(ii)** a posterior channel. **(B)** Exponent across Hori stages at **(i)** an anterior and **(ii)** a posterior channel, faint coloured lines indicate individual participants, black line indicated group mean, black bars indicate standard error of the mean. **(C)** same as **(B)** but for theta SNR. **(D)** same as **(B)** but for alpha SNR. Exponent units are a.u. Hz^-1^, and SNR units are a.u.

**Figure 7.**
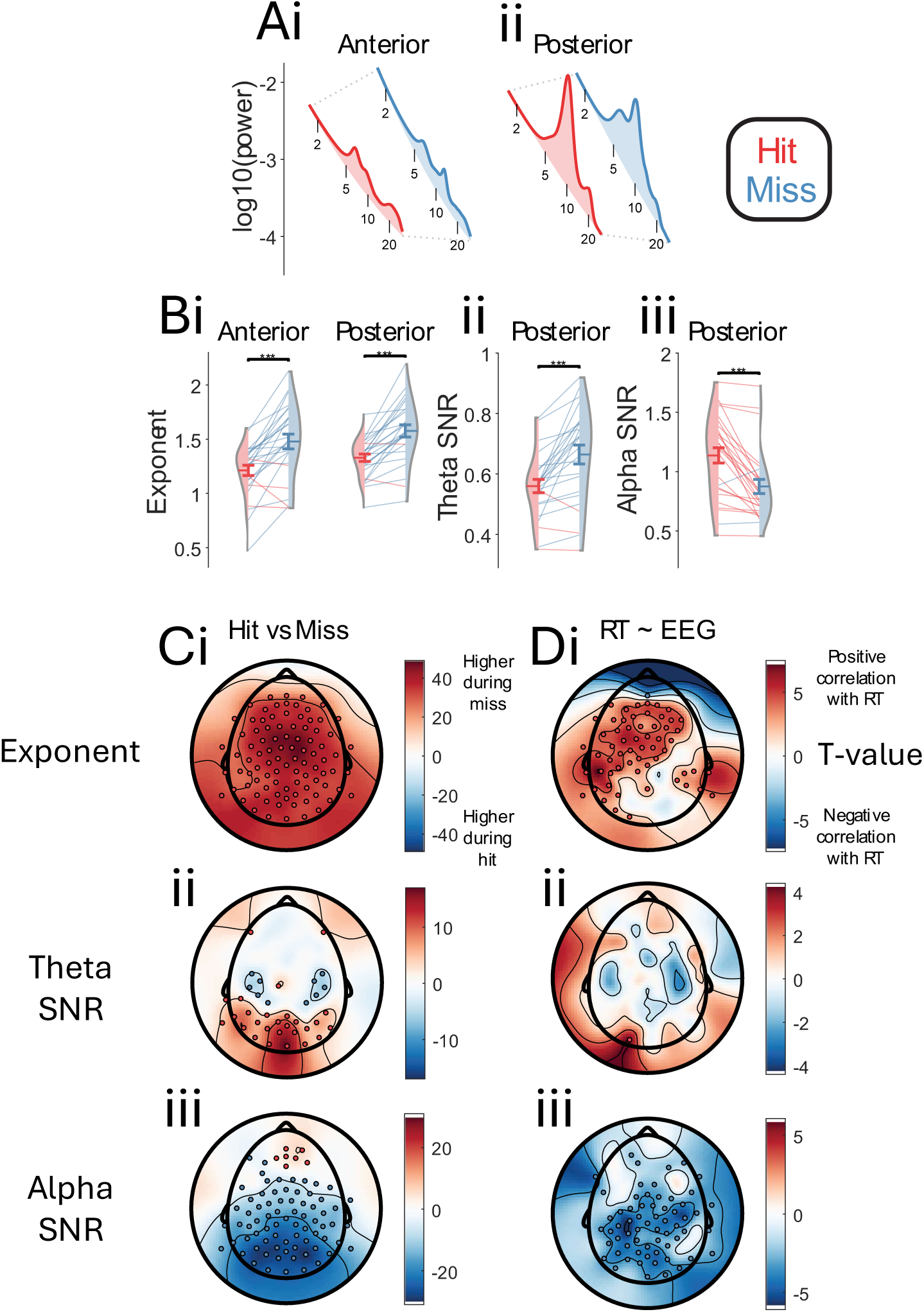
Relationship Between Aperiodic and Oscillatory Features and Responsiveness. **(A)** Power spectra for hits and misses at **(i)** an anterior and **(ii)** a posterior channel. **(B)** Exponent for hits and misses at **(i)** an anterior and a posterior channel, violins show the distribution in each condition, individual participants are represented by faint lines, the colour of which indicates which condition is higher (hit or miss), error bars indicate standard error of the mean, statistics marks indicate the results of mixed-effects models [exponent ∼ hit/miss + (1|participant)], *** *p<.001*. **(Bii)** same as **(Bi)** but for posterior theta SNR. **(Biii)** same as **(Bi)** but for posterior alpha SNR. **(C)** topographical representation of relationship between **(i)** exponent; **(ii)** theta SNR; **(iii)** alpha SNR in 4 seconds preceding auditory stimuli and response (hits vs miss). **(D)** topographical representation of relationship between **(i)** exponent; **(ii)** theta SNR; **(iii)** alpha SNR in 4 seconds preceding auditory stimuli and reaction time. Red scatter points on topographies indicate a (FDR-corrected statistically significant, p<.05) positive relationship between subjective sleepiness and the EEG feature in question, whilst blue scatter points indicate a (FDR-corrected statistically significant, p<.05) negative relationship. Exponent units are a.u. Hz^-1^, and SNR units are a.u.

In order to explore vigilance as a continuum within responsiveness, we related our EEG metrics of choice with reaction time, following the rationale that lower vigilance would bring about slower responses (**Figure 7D**). Consistent with expectations, a steeper exponent related to slower reactions (lower vigilance), with a predominantly anterior topography (**Figure 7Di**), whilst higher posterior alpha SNR related to faster reactions (higher vigilance) (**Figure 7Dii**). Theta SNR, however, showed a very minimal significant relationship with reaction time, confined to a single channel (**Figure 7Diii**).

#### Spectral Components Show Distinct but Symmetrical Trajectories Across the Loss and Recovery of Responsiveness

Finally, we turned our attention to transitions between states, that is, the moments just before and after responsiveness was lost or regained. To investigate this, we sorted the data according to its temporal distance from a transition. For example, if a hit was followed by 60 seconds of hits and then a miss, that epoch was assigned a value of –60 seconds on the green line (hit → miss) in **Figure 8**. Conversely, if a hit was preceded by a miss and 60 seconds of misses, it was assigned a value of +60 seconds on the purple line (miss → hit) in **Figure 8**. This approach allowed us to quantify both behaviour and physiology as a function of proximity to losing or regaining responsiveness.

**Figure 8.**
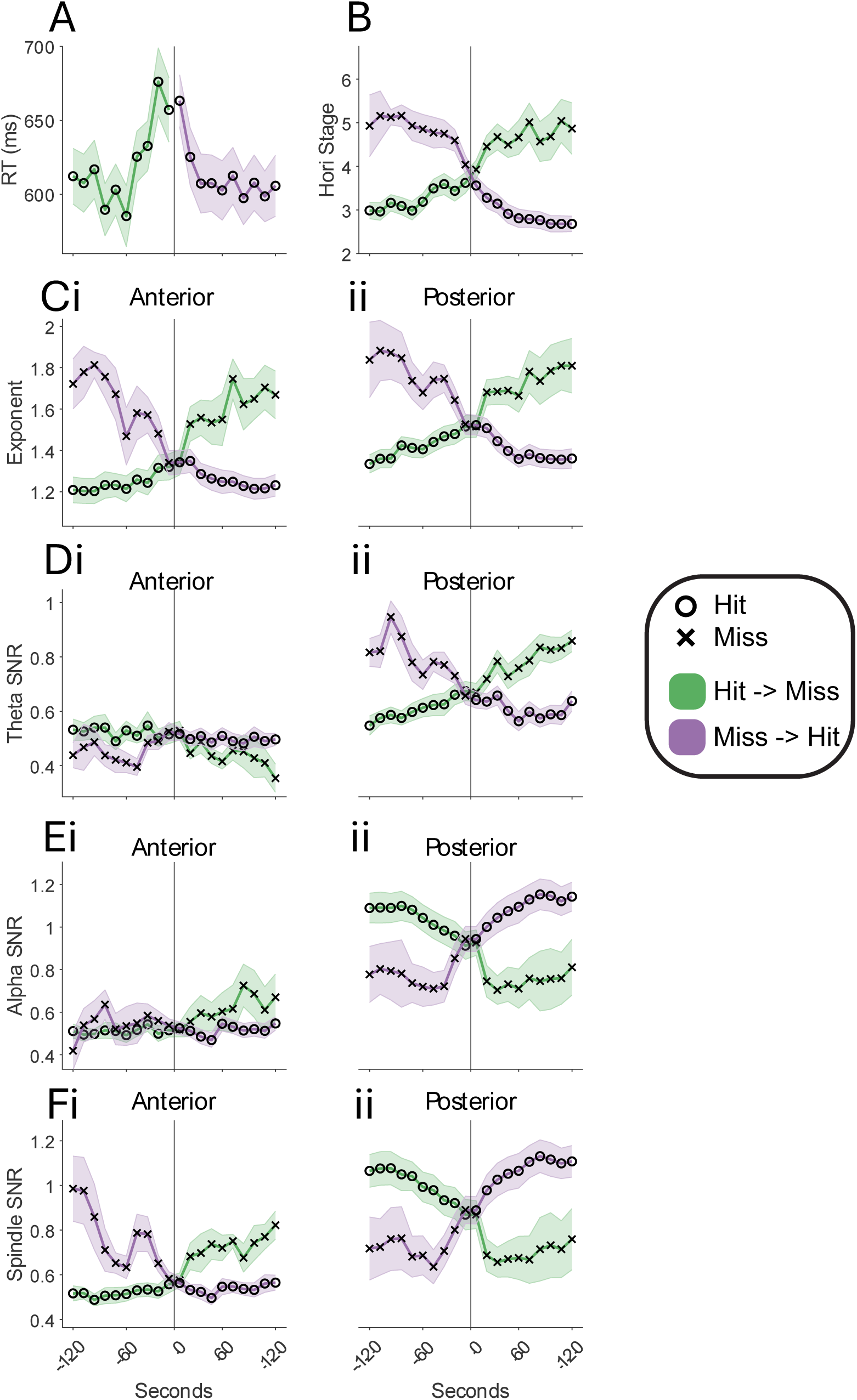
The trajectory of different features during transitions in responsiveness. Shown is the trajectory of a feature (e.g., anterior exponent) in the 2 minutes preceding and proceeding a transition in responsiveness. Green timeseries indicate a transition from hitting to missing (i.e., losing responsiveness), purple timeseries indicate a transition from missing to hitting (i.e., regaining responsiveness). ***O*** symbols indicate a hit, whilst ***X*** symbols indicate a miss. Plotted are **(A)** Reaction time, **(B)** Hori stage, **(C)** Exponent, **(D)** theta SNR, **(E)** alpha SNR, **(F)** spindle SNR. Shaded error bars indicate standard error of the mean. Note, for reaction time, the timeseries is discontinued when responsiveness is lost, since there is no reaction time to plot. Exponent units are a.u. Hz^-1^, and SNR units are a.u.

In line with the other portions of our investigation, we theorised that vigilance is not an *all-or-none phenomenon*, i.e., we would not expect a dramatic change in the EEG at the moment responsiveness changes nor would we expect strong consistency within a state, rather we anticipate a more gradual dynamic in the physiology around these transitions if vigilance indeed exists as a continuum. Similarly, we expected the speed of responsiveness itself to slow before it was lost entirely, and indeed reaction time showed a dynamic either side of these transitions, slowing (∼600 ms to ∼700 ms) in the 60 seconds before responsiveness was lost, and taking some moments to return to baseline after responsiveness returned (**Figure 8A**). Hori stage also saw a gradual increase in the minutes preceding a loss of responsiveness and this rather linear trend continued after the transition, whilst transitions in the opposite direction (i.e., regaining of responsiveness) showed an approximately mirrored dynamic (**Figure 8B**). Similarly linear and mirrored trajectories were observed for anterior exponent (**Figure 8Ci**), posterior exponent (**Figure 8Cii**), and posterior theta SNR (**Figure 8Ciii**). Posterior alpha SNR, which we have also repeatedly highlighted as an effectual readout of vigilance, showed a strikingly different dynamic, presenting a gradual drop in the ∼60 seconds preceding the transition from responsiveness, followed by a sharp drop immediately *after* the transition and no specific trajectory (i.e., flat) in the proceeding 2 minutes (**Figure 8Dii**). Despite its complexity, relative to the aforementioned features, the mirror image was again observed during the regaining of responsiveness. Anterior spindles, meanwhile, showed very little variance within responsiveness. This follows logically, since genuine *sleep* spindles should not be present during “wakefulness”. However, in the moments preceding a return of responsiveness in particular, a conspicuous non-linearity was observed in anterior spindles – a bump (deviation from linear trend) in the trajectory ∼30-40 seconds prior to transition. Though it is beyond the scope of this work, we suggest this might be capturing the infraslow fluctuation of sleep spindles (a non-invasive proxy of noradrenaline) which has strong implications for vigilance [62–64].

## 4 DISCUSSION

Here, we have leveraged high-density EEG and time-resolved spectral parameterisation to quantify the physiological substrates of vigilance with a greater fidelity than traditional sleep scoring or band power analyses. Whilst previous studies have largely focused on the comparison and contrast of somewhat arbitrarily defined discrete states, we have treated vigilance as a continuum, and expanded and substantiated this notion by implementing additional subjective and behavioural measures of vigilance.

Despite the longstanding use of neural oscillations to define vigilance, we show that the aperiodic component of the EEG power spectrum unequivocally outperforms its periodic counterpart, showing a striking consistency across definitions of vigilance, brain states, individuals, and datasets. The exponent emerged as a more robust, spatially consistent, and temporally linear marker of vigilance, operational both between- and within-states.

### Aperiodic and Periodic Components likely Reflect Distinct Neural Processes

Though periodic and aperiodic features both capture meaningful aspects of brain state, their differing spatio-temporal profiles indicate that they, importantly, may reflect distinct underlying systems. Aperiodic activity varied smoothly and globally with gradual changes in responsiveness, reaction time, and sleepiness. In contrast, oscillatory features, such as alpha, theta, and spindle-band activity, showed more complex, regionally specific, and often non-linear dynamics.

The precise biological underpinning of these distinct periodic and aperiodic profiles cannot be easily determined. Whilst there have long existed detailed and largely compatible hypotheses on the mechanisms of the brain’s oscillations [65–69], theoretical accounts of aperiodic neural activity remain ununified. The spectral exponent has been linked to a variety of mechanisms, including excitation/inhibition balance [39,53,55], synaptic timescales [53], criticality [70], and cortical bistability [54], but no single interpretation currently prevails. However, these theoretical models are not necessarily mutually exclusive [53], and indeed this multiplicity may be partially explained by the fact that the spectral exponent is a compound measure: sensitive to changes in both low- and high-frequency power, which may capture multiple processes.

Whilst we are cautious not to over-interpret at this nascent stage of aperiodic neuroscience, our data support the notion that the exponent might well capture *global* phenomena such as overall neuromodulatory tone [71] (i.e., excitation/inhibition balance) or gradual changes in the firing mode of thalamo-cortical neurons/interactions [72], and that oscillatory activity may better reflect *local* processes or circuits which undergo more frequent and abrupt changes.

### Utility

Whilst physiological interpretation might remain out of reach, this does not preclude utility. In terms of clinical evaluation of vigilance, the case for the exponent as a marker of vigilance is very clear. It captures vigilance independent of behavioural stage, meaning there is no requirement of time-consuming and labour-intensive sleep scoring for interpretation. This is not true of band-limited power, the meaning of which is highly state-dependent. For similar reasons, the exponent could provide a more holistic view of vigilance than the current clinical standards (MWT and MSLT) which depend on a binary classification of wakefulness and sleep. For instance, if two individuals maintain wakefulness throughout an MWT or MSLT, they will be given the same score, even if one is extremely drowsy and one is extremely alert, since neither fell asleep. We contend that these scenarios are quite different from each other, should be treated as such, and that the EEG’s aperiodic component may provide a more nuanced readout of vigilance than these tests. In addition, micro-sleep episodes are often not accounted for by these tests, but could potentially be captured by the spectral slope.

Finally, since the exponent does not suffer the spatial specificity of oscillatory activity, and can hence be recorded from anywhere on the scalp, the need for high-density EEG montages or technical expertise is reduced. Such measures are also easily applicable to low-density EEG systems in the clinic or home setting. For these reasons, it would be valuable to compare the spectral slope against traditional readouts of MSLT and MWT recording and behavioural tests.

Our findings challenge the longstanding assumption that oscillatory activity, particularly the fading out of alpha oscillations, is an appropriate marker of diminishing vigilance. While alpha did exhibit expected patterns, such as sharp decreases during loss of responsiveness, it was less predictive overall than the spectral exponent and more sensitive to spatial variation. Oscillations may still be valuable for detecting discrete state shifts, but their irregular dynamics limit their utility for continuous tracking. This is of especial importance, considering that clinical scoring (on which the MWT and MSLT are based), is highly dependent on the drop out of alpha activity.

Studying the aperiodic component of neural spectra also side-steps a number of issues associated with traditional frequency-limited approaches, requiring less arbitrary discretisation of the spectrum, potentially showing a consistent meaning across states, remaining operational in the absence of oscillations, and insensitive to frequency deviations between individuals. Far fewer parameters are required to be set, and hence the simplicity of the measure is superior. Nevertheless, the approach based on the spectral slope still relies on the Fourier transform – a technique fundamentally poorly-suited to EEG signals, which violate many of its core assumptions. It addresses many issues with oscillation-centric analysis, but rather avoids the fundamental one – in the end we may have to face this issue, working towards better methods to operationalise meaning in the EEG.

### Cautions, Limitations, and Future Directions

Despite the coherent and convincing nature of the results presented here, there are several important limitations which warrant attention.

Firstly, while we found consistent effects across three datasets, replication in larger, more diverse, and clinical populations is needed. Those features underpinning vigilance in healthy populations may not generalise to *pathological* imbalances of vigilance.

Secondly, we have only investigated wakefulness and NREM sleep here, and have not expanded our investigation to include REM sleep, which may follow very different trajectories [45,47]. However, in the use cases we have put forward (i.e., excessive or insufficient vigilance), the transitions of interest are considerably more likely to be between wakefulness and NREM sleep, and hence our findings remain practical.

Thirdly, although the EEG provides valuable information, other physiological measures - such as pupil diameter, heart rate variability, respiration, muscle tone, and body movement - may also provide unique insights into vigilance. Integrating these modalities could offer a more complete picture.

Finally, our analysis did not distinguish whether changes in spectral slope arose from high-amplitude low-frequency events (such as K-complexes or slow waves [31,61,73,74]) or from shifts in higher-frequency activity. In addition, although the main goal of this work was to simply compare how different spectral components capture vigilance, other *event-based* measures (e.g., slow wave detection) may also prove effective and insightful.

Future research would do to well to expand the current work by assessing the sensitivity of aperiodic features to pharmacological and behavioural interventions, examining their stability across populations and clinical contexts, exploring integration with other physiological signals, and further defining the origins of changes to neural power spectra.

## Conclusion

Our findings provide strong evidence that the spectral exponent - a feature of aperiodic EEG activity - offers a simpler, more scalable marker of both subjective and objective aspects vigilance than traditionally implemented measures of neural oscillations. Its consistent, global, and gradual dynamics make it better suited than oscillatory activity for capturing brain state across time. While both aperiodic and periodic components offer insight, their distinct profiles underscore the need to move beyond an oscillation-centric model of vigilance. The spectral exponent represents a promising foundation for advancing both basic research and clinical applications involving consciousness, sleep, and arousal, as well as providing a potentially attractive target for characterisation and treatment of disorders associated with disturbed vigilance.

## 5 METHODS AND MATERIALS

### 5.1 Participants

Studies 1 and 2 comprised the same 18 healthy participants (age 37.7 ± 7.9 years (mean ± SD), range: 25 – 51, 12 females) with regular:

- Bed and rise times
- Subjective sleep quality (Pittsburgh Sleep Quality Index [75] < 5)
- Chronotypes (Horne and Ostberg morningness-eveningness [76] <70 and >30)
- Daytime sleepiness (Epworth Sleepiness Scale [77] < 10)

They were free from all medical, neurological, and psychiatric disorders affecting sleep, did not take regular medicine (aside from birth control) and were not pregnant. Data from these participants were also used in these previous studies: [31,57].

Study 3 comprised 29 right handed healthy participants, with a *high* daytime sleepiness (Epworth Sleepiness Scale [77] >7), who were asked not to consume stimulants (e.g., caffeine) in the hours before the experiment. Data from this study was used in this previous study [20], before being made only available online (https://doi.org/10.17863/CAM.18707-dataset2).

### 5.2 Study Designs

Study 1 comprised the first hour of an undisturbed nocturnal sleep episode taken from the baseline night, beginning at lights off – approximately 23:00.

Study 2 comprised a serial awakening protocol [56]. Participants underwent two experimental nights, at least one week apart, in which they were repeatedly awakened throughout the night and, immediately upon awakening, interviewed about their subjective experience. Awakenings were conducted using a 1.5 s computerised alarm sound by a researcher monitoring the participant via video and EEG from a control room. Awakenings from NREM were carried out ∼10 minutes after the first sleep spindle. Only awakenings from NREM sleep (N2 awakenings (11.1 ± 4.3 awakenings (mean ± SD), range 6-17), N3 awakenings (9.7 ± 5.6 awakenings (mean ± SD), range 1-22), total NREM awakenings (20.8 ± 4.4 awakenings (mean ± SD), range 14-29)), with N2 and N3 sleep being combined, and only the question on subjective sleepiness (1-5) were analysed in the present work – but see [31] for full experimental details.

Study 3 comprised an auditory masking task, in which participants were required to press a button when a target sound was administered. The sound was a beep randomly masked by different noise durations, so as to be slightly difficult to detect without sufficient attention. The experiment was conducted in a dark room, whilst the participant reclined in a comfortable chair in the dark with their eyes closed, and were permitted to fall asleep. An average of 543 trials (SD 62.8, range 402-631) were administered, and the experiment lasted a maximum of 120 minutes.

### 5.3 EEG acquisition, scoring, and re-referencing

EEG data in studies 1 and 2 were recorded at a sampling rate of 500 Hz using a 256-channel net (Electrical Geodesics Inc), on-line referenced to the vertex (Cz). See [31] for full pre-processing. EEG data were scored in 30-second epochs, according to the AASM conventions [21].

EEG data in study 3 were recorded at a sampling rate of 500 Hz using a 128-channel net (Electrical Geodesics Inc), on-line referenced to the vertex (Cz). Peripheral channels were removed by the original authors, leaving only those 93 electrodes on the scalp. See [20] for full pre-processing. EEG data were scored in 4 second epochs, according to the Hori conventions [22,23].

In all cases, EEG data were subject to a laplacian-type re-referencing (Scalp Current Density using FieldTrip [78]), in which each channel is referenced to its immediate neighbours.

### 5.4 Parameterisation of Power Spectra

Power spectra were computed and parameterised using the SPRiNT [36] extension of the BrainStorm toolbox. Spectra were computed 1.5-30 Hz, using 3-second windows in studies 1 and 2 and 4-second windows in study 3 (to account for 4-second trial length), with a 50% overlap, and averaged over two sliding windows. Spectra were parameterised according to the following parameters across studies: aperiodic-mode: *fixed*; peaks: *gaussian*, maximum peaks: *4*; minimum peak height: *0.c*; minimum peak width *1.5*; maximum peak width *c*; peak threshold: *2* (SDs).

At each channel, the spectral exponent was extracted, along with the SNR of all detected oscillatory peaks. SNR values were sorted on the basis of canonical frequency bands; theta (4-8 Hz), alpha (8-12 Hz), and sigma (9-16 Hz). Note: whilst the exponent should be negative (i.e., an exponential decay of power with increasing frequency), the SPRiNT toolbox expresses it as a positive value. In general, we also find this to be more intuitive when describing/discussing these data – with a higher exponent indicating a steeper slope.

### 5.5 Regions of interest

In many cases, analyses are restricted to smaller, anterior and posterior, clusters of midline channels. The anterior cluster comprised Fz, its immediately anterior neighbour, and its immediate posterior neighbour. The posterior cluster comprised Oz, its immediately anterior neighbour, and its immediate posterior neighbour.

### 5.6 Statistical Analysis

Linear mixed effects models were used for all statistical analyses, with participant as a random effect. In the case of topographical analyses, FDR corrections were conducted to control for multiple comparisons.

### 5.7 Aspects of Vigilance

The measures of interest were as follows:

Study 1: AASM sleep stage, time in bed, time in NREM sleep; study 2: time from awakening, subjective sleepiness; study 3: Hori stage, responsiveness (i.e., hit or miss); reaction time, time from transition (between responsiveness and unresponsiveness in each direction).

#### CRediT

**HH**: conceptualisation, software, formal analysis, data curation, writing – original draft, writing – review and editing, visualisation; **AMS**: investigation, resources, data curation, writing – review and editing; **JC**: investigation, resources, data curation; **FS**: resources, writing – review and editing, supervision, funding acquisition.

